# Impact of a DNase I and Vancomycin Drug Combination against a Mature Methicillin Resistant *Staphylococcus aureus* (MRSA) *in vitro* Biofilm

**DOI:** 10.64898/2026.02.04.703709

**Authors:** Rex C. Moore, Hugh D.C. Smyth

## Abstract

Antimicrobial resistant infections present a concerning and expanding healthcare problem that is compounded by the reduction of antimicrobial research by the pharmaceutical industry. Additionally, the currently used antimicrobials consistently present less than ideal clinical treatment outcomes. This contention is supported by *in vitro* analysis in appropriate models. Here, in mature methicillin resistant *Staphylococcus aureus* (MRSA) *in vitro* cultures, we tested multiple antimicrobials and showed that for a biofilm grown for 72 hours, no antimicrobial tested was able to completely eradicate the biofilm even after 24 hours of exposure. However, the addition of an enzymatic biofilm-dispersal agent (DNase I or Proteinase K), greatly improved the performance of vancomycin and tigecycline in this *in vitro* model. Despite the improved performance in the presence of the dispersal agent, a high concentration of antimicrobial, 2000 µg/mL, was needed to completely eradicate the infection as demonstrated by analyses using both a traditional XTT assay as well as a subculture assay to account for persister cells. It was shown that the addition of DNase I improved the diffusion of vancomycin through the biofilm. This suggests vancomycin efficacy is limited by the biofilm. The presented work provides a potential avenue for future treatments of MRSA lung infections by utilizing a traditional antibiotic combined with a passive dispersive agent.

## 1.2 Introduction

A continuing problem that impacts healthcare in the US and around the world is the emergence and prevalence of antimicrobial resistant (AMR) infections. It is estimated that AMR infections were responsible for more than six million deaths globally in a single year with the expected death toll to balloon to 10 million by 2050 (1, 2). The pathogens that are responsible for these infections can vary depending on the region of the world, but one of particular concern is methicillin resistant *Staphylococcus aureus* (MRSA). A CDC report from 2019 found that within the US, MRSA infections accounted for an estimated $1.7 billion in attributable healthcare costs; however, the impact of MRSA infections goes far beyond with more than 323,000 cases in hospitalized patients leading to more than 10,000 deaths in a single year (3, 4). Nosocomial pneumonia infections, which are defined as a pneumonia infection that occurs 48 hours or more after hospital admission (5), are particularly hard to treat. It’s estimated that 15% of nosocomial infections can be attributed to pneumonia which results in longer hospital stays, increased treatment costs, and a threefold increase in mortality (6–8). MRSA is believed to be responsible for between 20-40% of all hospital-acquired pneumonia and ventilator-associated pneumonia infections (6, 7).

The therapeutic treatment options available for MRSA infections have generally limited clinical success and at times high mortality rates ranging from 5-60% depending on the site of infection and patient factors such as age, organ dysfunction, or ICU status (8, 9). Two of the most commonly used antibiotics for MRSA lung infections are linezolid and vancomycin (10–12). Despite the vastly different physicochemical properties and pharmacokinetic profiles, the two antibiotics display similar clinical cure rates. The definition of clinical cure rate can vary depending on study criteria, and despite the use of linezolid and vancomycin globally, there is still considerable debate about whether one is superior to the other (8, 13, 14). Regardless of the selection of a first line agent, improvements in treatment options are not only warranted but needed (15, 16). One of the biggest issues with treating established infections with biofilm involvement is achieving a sufficiently high drug concentration at the site of infection (17, 18). The conundrum for systemically administered antibiotics are dose limited due to side effects (17, 19, 20). Where appropriate, various routes of localized delivery have been investigated with varying degrees of success. For example, for pulmonary infections, marketing approval of tobramycin inhalation solution, TOBI® Podhaler®, Aztreonam for inhalation solution, colistin solution and dry powder, and amikacin liposome inhalation suspension illustrate the viability of this approach (albeit primarily in gram negative pathogens highlighting the unmet need for MRSA and other gram-positive pathogens). In contrast, several inhaled antibiotics in late-stage development have also failed to achieve marketing approval including vancomycin (AeroVanc™ developed by Savara Pharmaceuticals (21)), ciprofloxacin (BAYQ3939 developed by Bayer (22), and Pulmaquin® developed by Aradigm Corp (23)).

MRSA infections, especially in the lung, are difficult to treat due to the ability of the pathogen to form biofilms (18, 24–26). Biofilms are complex heterogeneous communities that consist of an extracellular matrix containing proteins, extracellular DNA (eDNA), polysaccharides and other components (27–29). The three-dimensional structure provides a complex defense mechanism that protects the bacterial cells from environmental stresses, the host immune response, and antimicrobial agents (29).

Additionally, the presence of a biofilm creates nutrient gradients and water channels which leads to the development of persister cells (30, 31). Persister cells are essentially dormant or slow growing cells that are incredibly difficult to treat with antimicrobials due to the limited metabolic activity (30, 32). Biofilm colonies also facilitate horizontal gene transfer which can lead to increased tolerance and resistance mechanisms. The difficulty in treating the complex biofilms has led to the pursuit of new agents and treatment methods (27).

The purpose of this work was to identify potential local treatment improvements using existing therapeutic agents with or without biofilm dispersal enzymes in an *in vitro* model of mature MRSA biofilms. First, biofilm growth conditions (growth media and time for biofilm growth) were studied to establish a robust biofilm model system. Second, determination of the minimum inhibition concentration (MIC) was utilized as a screening tool for six antimicrobials in this model. Following drug screening, assessment of the minimum biofilm eradication concentration (MBEC) was performed to determine the appropriate antimicrobial and treatment conditions with or without the addition of biofilm dispersive agents in the form of Proteinase K and DNase I. Identification of successful treatment conditions was determined using an XTT assay as well as a broth subculture method to account for slow growing persister cells. This work will help improve the knowledge needed to treat mature MRSA biofilm infections using existing antimicrobials as well as combination treatment regimens.

## 1.3 Results

### 1.3.1 An *in vitro* Model of Mature Biofilms: Biofilm Biomass

Before the antimicrobial susceptibility of the two MRSA (ATCC 33592 and BAA - 41) biofilms could be tested, an *in vitro* model of mature biofilms that demonstrated consistent, strong, and mature biofilms was needed. Consistent growth was defined as measured growth with relative standard deviations of 15% or less. A strong biofilm was defined using the classification systems proposed by Singh et al. Briefly, a strong biofilm occurs when the absorbance value of the biofilm is 4-fold or greater than the absorbance of the negative control (33). To identify biofilm conditions yielding consistent and strong biofilms, multiple growth medias and conditions were identified from literature and assessed using a 0.5% crystal violet biomass assay at different time points (29, 33, 34). The biofilms were formed on flat-bottom, cell treated polystyrene plates for a period of 1, 3 or 5 days of growth in six different growth medias. The growth medias, tryptic soy broth (TSB) and brain heart infusion broth (BHI), have been commonly used for MRSA strains. Additionally, the growth media was supplemented with either glucose or sodium chloride to determine how changes in growth media composition might improve the biofilm biomass. As shown in Figure 1, similar biofilm growth characteristics were seen for both stains in BHI supplemented with 1% glucose grown for 3 days. These conditions resulted in a larger (i.e. stronger) and consistent biofilm, evident by the relative standard deviation being less than 15%. MRSA strain 33592 produced biofilm of greater biomass, evidenced by the increased absorbance value, than MRSA strain BAA-41. It is interesting to note that addition of NaCl to BHI resulted in a very low biomass for both strains while the addition of NaCl to TSB did not see a similar effect. The statistical analysis showed that both the effect of growth media (p-value < 0.0001) and incubation time (p-value 0.004) were significant in the resulting biofilm biomass. At the three-day timepoint, the comparison between BAA-41 BHI + 1% glucose compared to 33592 BHI + 1% glucose did not result in statistical significance. For future studies, BHI + 1% glucose was chosen as the growth media because of the strong and consistent biofilm that was formed.

**Figure 1.**
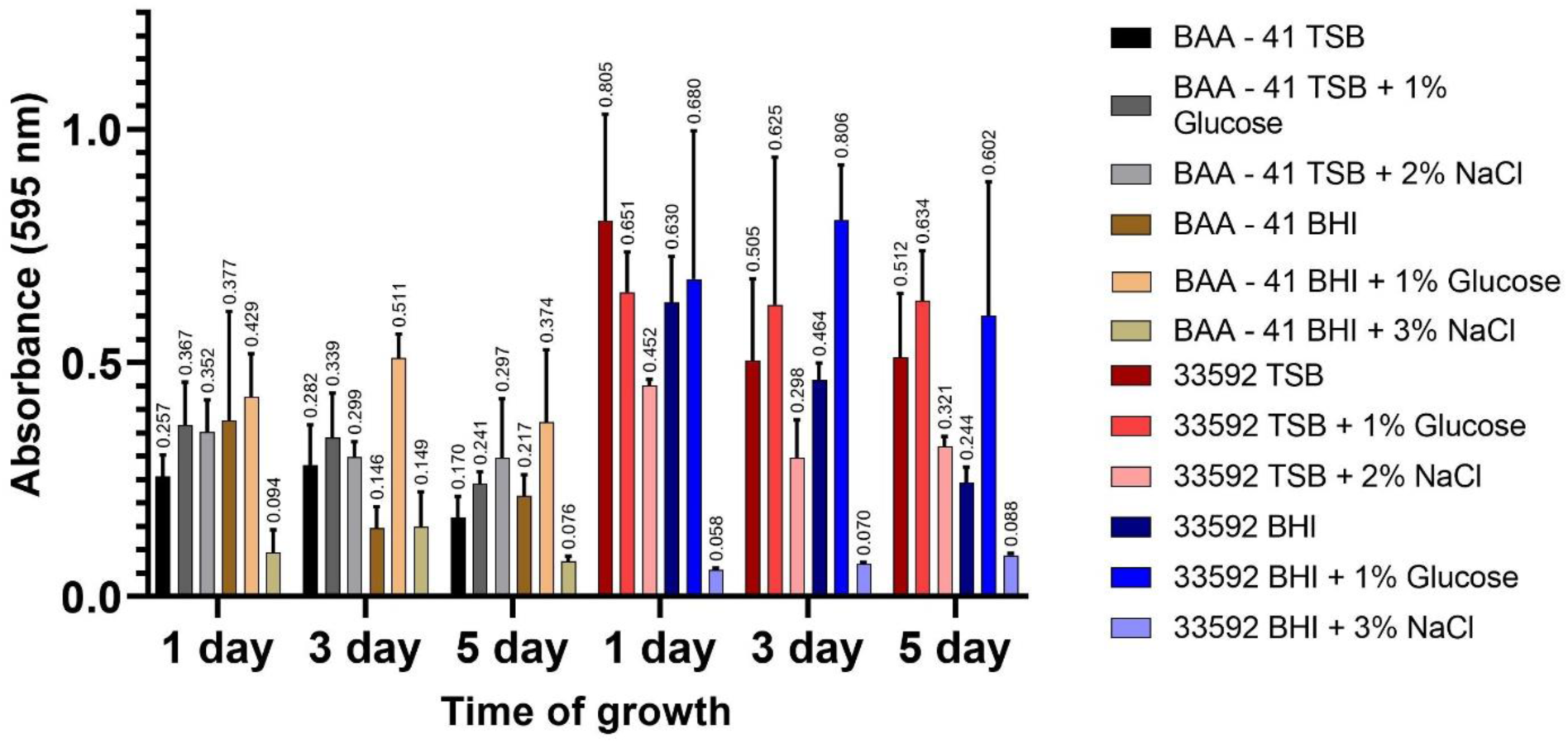
Biofilm biomass of two MRSA strains (ATCC #: BAA-41 and 33592) under different culture conditions. MRSA strains were incubated for 1-5 days at 37°C. Y-axis represents the mean and standard deviation of the biomass measured using 0.5% crystal violet at an absorbance of 595 nm. TSB: tryptic soy broth; BHI: brain heart infusion broth

### 1.3.2 An *in vitro* Model of Mature Biofilms: Colony Forming Units (CFUs)

In addition to biofilm biomass, the number of colony forming units (CFUs) for the biofilms grown under the different conditions was tested. The combination of biofilm biomass and CFU counts helps provide a more holistic understanding of the strength of a biofilm. In contrast to the biofilm biomass observations, fewer differences in CFU counts between the two strains of MRSA were observed (Figure 2). At the 3-day timepoint, BHI + 1% glucose media did not result in statistically different number of CFUs for the 33592 strain when compared to the other medias at the same timepoint. However, for the BAA-41 strain, there was a significant difference between the BHI and BHI + 1 % glucose media. It was determined that the selected growth media and incubation time were significant factors (P < 0.0001) based on the 2-way ANOVA statistical analysis that was completed.

**Figure 2.**
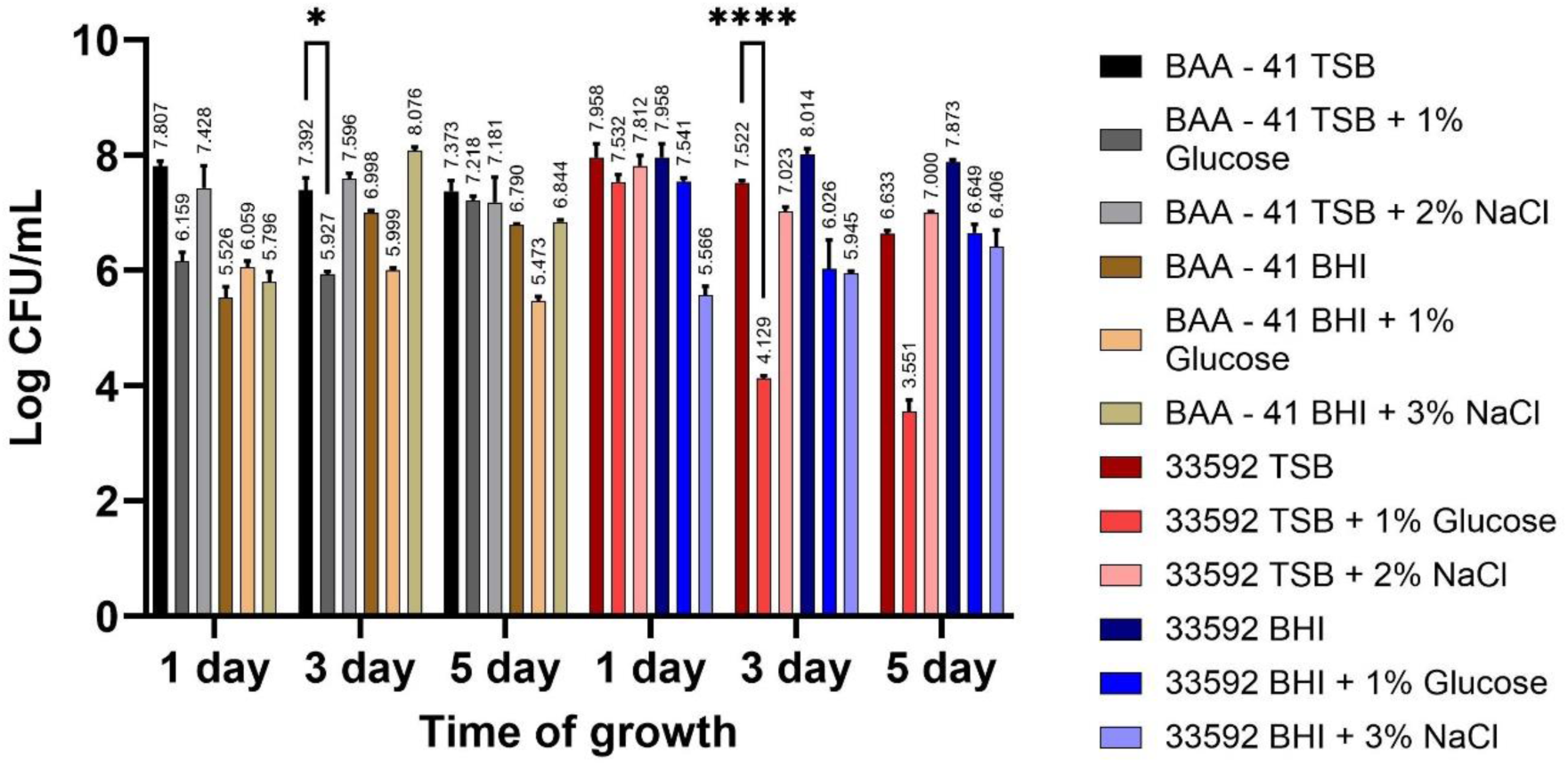
Log CFU/mL of two MRSA strains (ATCC #: BAA-41 and 33592) under different culture conditions. MRSA strains were incubated for 1-5 days at 37°C. Y-axis represents the mean and standard deviation of Log CFU/mL tested using a drop plate method.

### 1.3.3 Screening of Candidate Antibiotics: Minimum inhibitory concentration (MIC)

Following the determination of the best growth conditions for a robust, mature biofilm, minimum inhibitory concentration (MIC) testing was used to screen potential antibiotics to select candidates for further evaluation. A total of six antibiotics were chosen for this study. Three antibiotics (vancomycin, linezolid, and clindamycin) were identified from clinical treatment guidelines for MRSA pneumonia infections (10, 35). An additional three were chosen based on reports of *in vitro* and clinical success (36–38). Although MIC testing has its limitations, especially to measure the antimicrobial susceptibility when biofilms are present (39), MIC can provide valuable data to be used as a screening tool to determine potential resistance. As can be seen in Table 1, clindamycin, tobramycin, and niclosamide did not perform well with very high MIC values; however, vancomycin, linezolid, and tigecycline performed well and fell within the margin of error for accepted clinical breakpoints per CLSI guidelines (40).

**Table 1.** MIC for six different antimicrobials against two strains of MRSA.

| Drug (µg/mL) | BAA-41 | 33592 |
| --- | --- | --- |
| Clindamycin | >64 | >64 |
| Tobramycin | >64 | 32 |
| Niclosamide | >64 | >64 |
| Vancomycin | 4 | 4 |
| Linezolid | 4 | 2 |
| Tigecycline | <0.5 | <0.5 |

### 1.3.4 Biofilm Eradication Using Antibiotics and Dispersal Enzymes

Due to the limitations in the use of MIC testing for antimicrobial susceptibility of biofilm infections, minimum biofilm eradication concentration (MBEC) assays were performed (39, 41). A metabolic XTT assay was used to determine the viability of the biofilm following antibiotic exposure. The antimicrobial performance was determined based on the percent reduction in metabolic activity by comparing the treated biofilms to a normalized untreated biofilm control grown under the same conditions. Additionally, sterile media, under the same conditions as biofilm samples, were used to ensure sterility of the samples. Antibiotic exposure times varied from 6 to 72 hours with antibiotics added once at the beginning of the timepoint or replaced every 12 hours to simulate a twice daily dosing regimen. In Figure 3(a) and 3(b), it can be seen that no antibiotic by itself was able to completely eradicate the biofilm viability. It is interesting to note that for BAA-41 biofilms, vancomycin appears to be the best performing antimicrobial followed by linezolid and tigecycline (Figure 3(a)) but based on a two-way ANOVA analysis none of the antimicrobials demonstrated statistically different performance over the concentrations and exposure times tested. In the case of 33592, the stronger biofilm model, tigecycline appeared to perform better with the largest reduction in metabolic activity followed by linezolid and finally vancomycin (Figure 3(b)). Statistical analysis showed that a difference in the performance of vancomycin and tigecycline occurred at concentrations of 500 and 750 µg/mL when antimicrobial exposure times of 12 hours were used. Additionally, linezolid and tigecycline were statistically different at concentrations of 500 and 2000 µg/mL when exposed for 24 hours. Despite the very high concentrations tested, higher than what is typically observed clinically after systemic administration, the antimicrobial treated biofilms still showed metabolic activity, and bacterial cells were still viable following subsequent culture.

**Figure 3.**
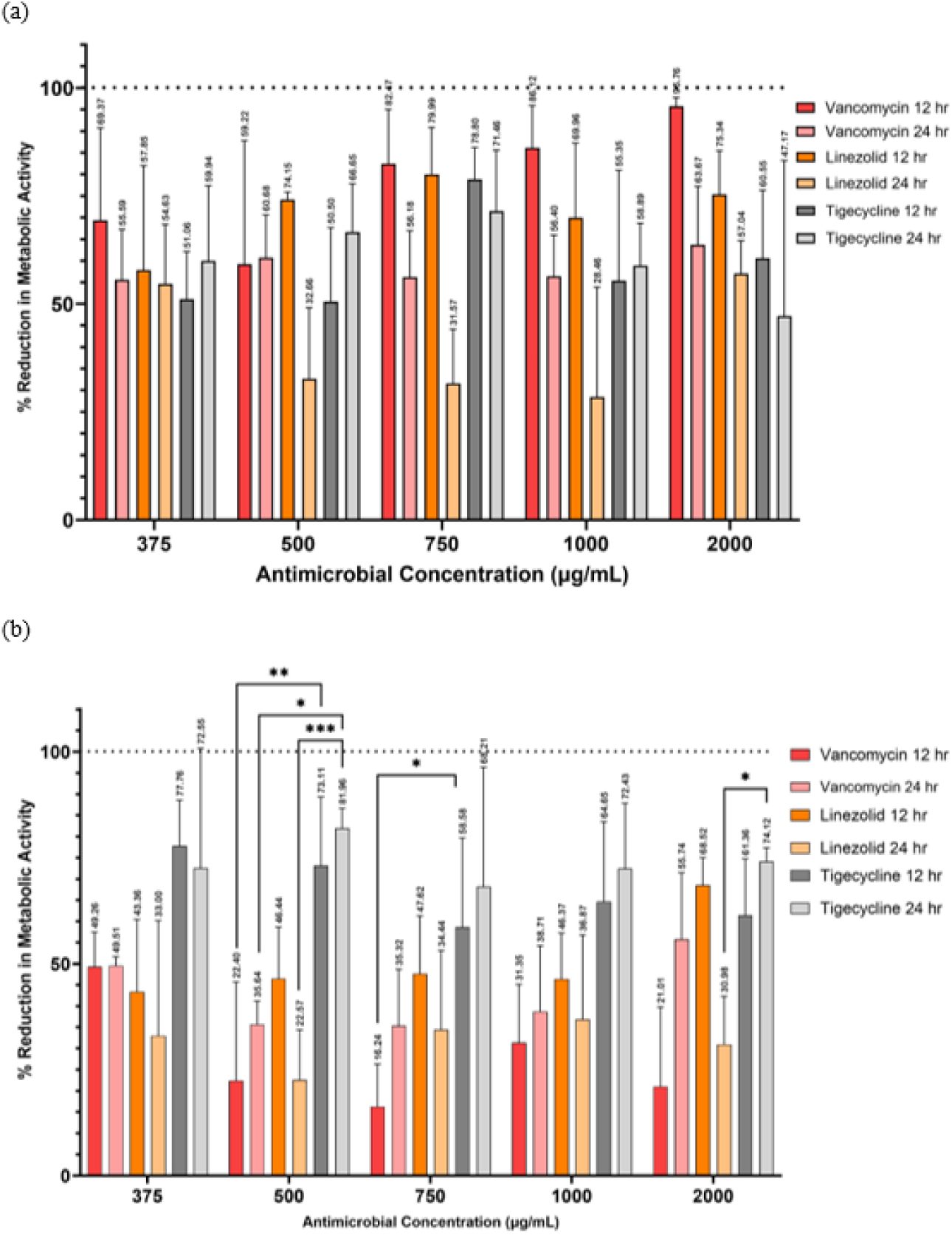
Antimicrobial susceptibility of MRSA strains BAA-41 (a) and 33592 (b). Both strains of MRSA biofilm were grown for 72 hours in BHI broth supplemented with 1% glucose before antibiotic exposure ranging from 12 hours to 24 hours. Antimicrobial susceptibility was determined based on reduction of metabolic activity determined with an XTT assay. Following analysis using a two-way ANOVA with Tukey method for multiple comparison, it was found that for strain BAA-41 there was no significance when comparing similar conditions while the significance for strain 33592 is designated by the asterisks (*P ≤ 0.05, **P ≤ 0.01, ***P ≤ 0.001).

Following the observation that no antimicrobial alone was able to eradicate the biofilm, vancomycin and tigecycline were combined with a biofilm dispersal enzyme (DNase I or Proteinase K (Pro-k)). The inclusion of DNase I (250 µg/mL) and Pro-k (100 µg/mL) was hypothesized to assist the performance of the antimicrobial by affecting the extracellular DNA or protein components of the biofilm structure that have numerous roles in the regulation of biofilms (27, 32, 42). The impact provided by the combination of the dispersal agents and tigecycline can be seen in Figure 4(a) and 4(b). Based on the XTT data, some of the drug-enzyme combination treatments were able to reduce the metabolic activity by 100%, achieving the goal of eradication, by XTT analysis. The combination of DNase I and tigecycline improved the antimicrobial performance by 10-40% (depending on the exposure time) for strain BAA-41 as seen in Figure 4(a). Improvements were also seen for strain 33592 (Figure 4(b)) albeit not quite as dramatic as strain BAA-41. The antimicrobial performance of vancomycin supplemented with DNase I or Pro-k resulted in a similar bactericidal improvement for strain BAA-41 as seen in Figure 5(a). Figure 5(b) shows a similar effectiveness of the vancomycin and dispersal agent combinations against the 33592 strain over a 72-hour period instead of the 24 hours that were required for the BAA-41 strain. It is important to note that vancomycin by itself was able to reach the 100% reduction in metabolic activity target threshold after a 72-hour period when the antibiotic was replaced every 12 hours as seen in Figure 5(a). An additional interesting observation was seen when the biofilm was pre-treated with DNase for 6 hours followed by 12 hours of vancomycin which resulted in a reduced antimicrobial performance, by about 20-25%, compared to the co-treatment combination of DNase and vancomycin for 12 hours (Figure 5(b)). This finding suggests that the drug-dispersal agent combination must be delivered at the same time for the improved antimicrobial performance. The staggered dose did result in a slight improvement in antimicrobial performance, albeit without statistical difference, compared to a single vancomycin alone dose over a similar exposure time for the lowest concentration shown, 750 µg/mL. For both strains, BAA-41 and 33592, neither dispersal agent, administered alone, was unable to kill the biofilm (data not shown).

**Figure 4.**
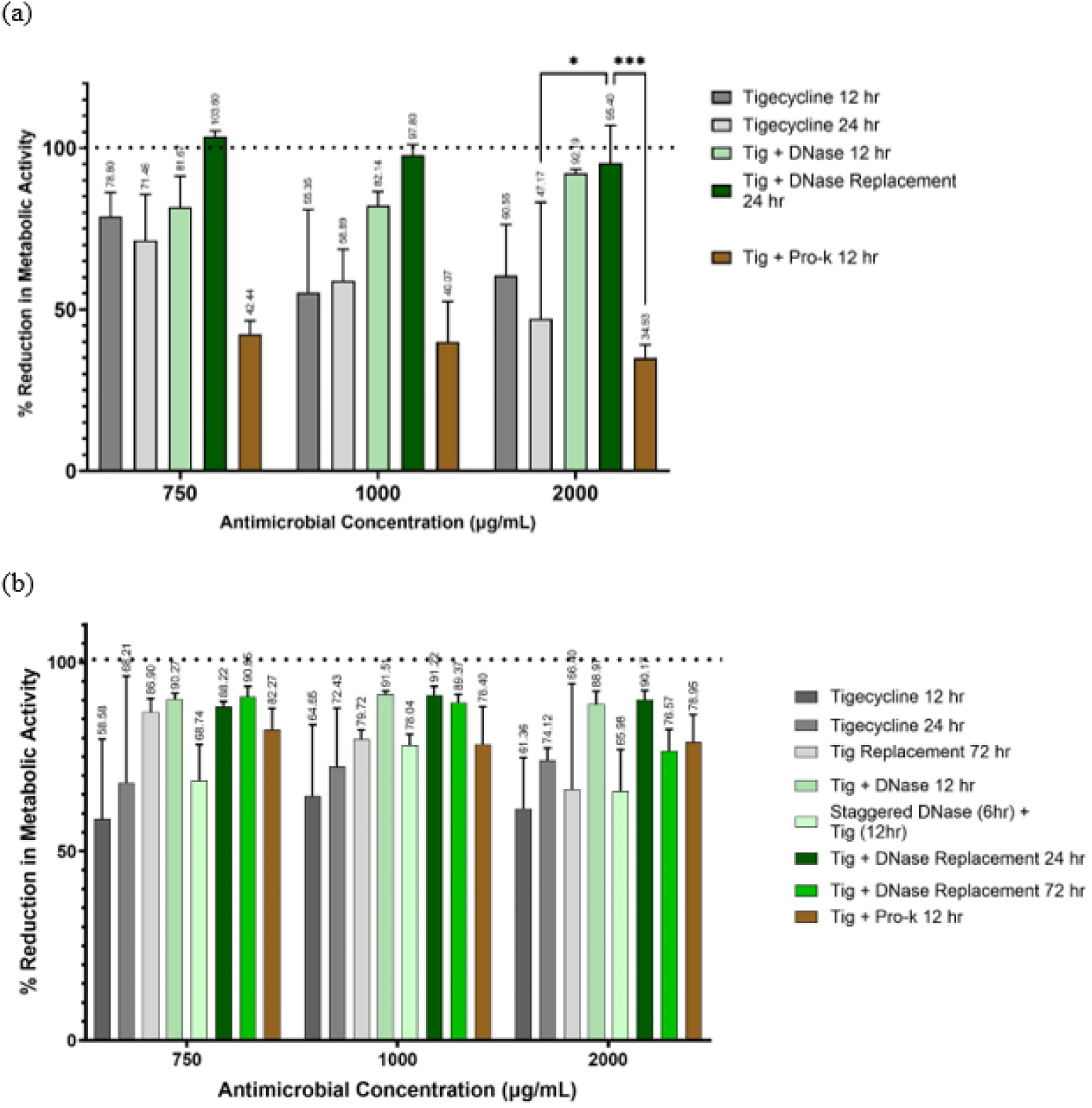
Antimicrobial susceptibility of MRSA strains BAA-41 (a) and 33592 (b) against combination therapy containing tigecycline and passive dispersal agents in the form of DNase I replacement in the therapy name in the legend to the right of each figure. Antimicrobial susceptibility was determined based on reduction of metabolic activity determined with an XTT assay. For strain BAA-41 there was significance, determined by a two-way ANOVA, denoted by asterisks (*P ≤ 0.05, **P ≤ 0.01, ***P ≤ 0.001, ****P ≤ 0.0001). There was no significance for strain 33592.

**Figure 5.**
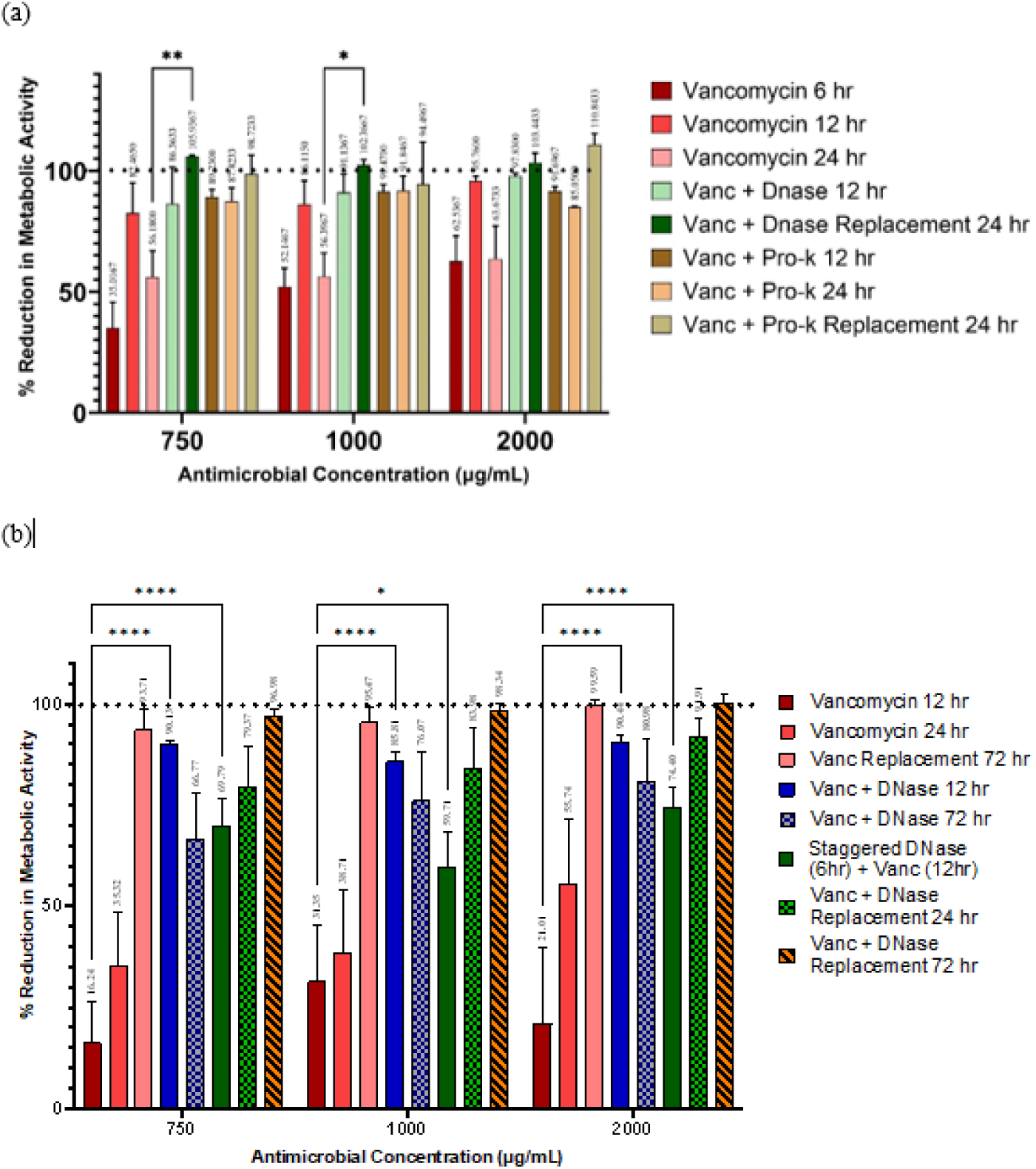
Antimicrobial susceptibility of MRSA strains BAA-41 (a) and 33592 (b) against combination therapy containing vancomycin and passive dispersal agents in the form of DNase I replacement in the therapy name in the legend to the right of each figure. Antimicrobial susceptibility was determined based on reduction of metabolic activity determined with an XTT assay. Significance determined by a two-way ANOVA is denoted by asterisks for both MRSA strains tested (*P ≤ 0.05, **P ≤ 0.01, ***P ≤ 0.001, ****P ≤ 0.0001).

Following the promising antibiofilm observations of the vancomycin and DNase co-treatment combination, various ratios of the two components were further tested. The concentration of vancomycin was ranged from 375 to 2000 µg/mL while the concentration of DNase was ranged from 125 to 1000 µg/mL. For strain BAA-41, as seen in Figure 6(a), the biofilm was treated with the combination for 24 hours where the treatment was replaced after 12 hours. The statistical analysis showed that the concentration of vancomycin did not have a significant effect on reduction in metabolic activity; DNase concentration, on the other hand, was found to be significant in certain conditions (Figure 6(a)). The impact of varied concentrations of the drug enzyme combination for strain 33592 can be seen in Figure 6(b). The limited difference in percent reduction of metabolic activity across the different concentrations tested can be seen in the figure which supports the lack of statistical significance found.

**Figure 6.**
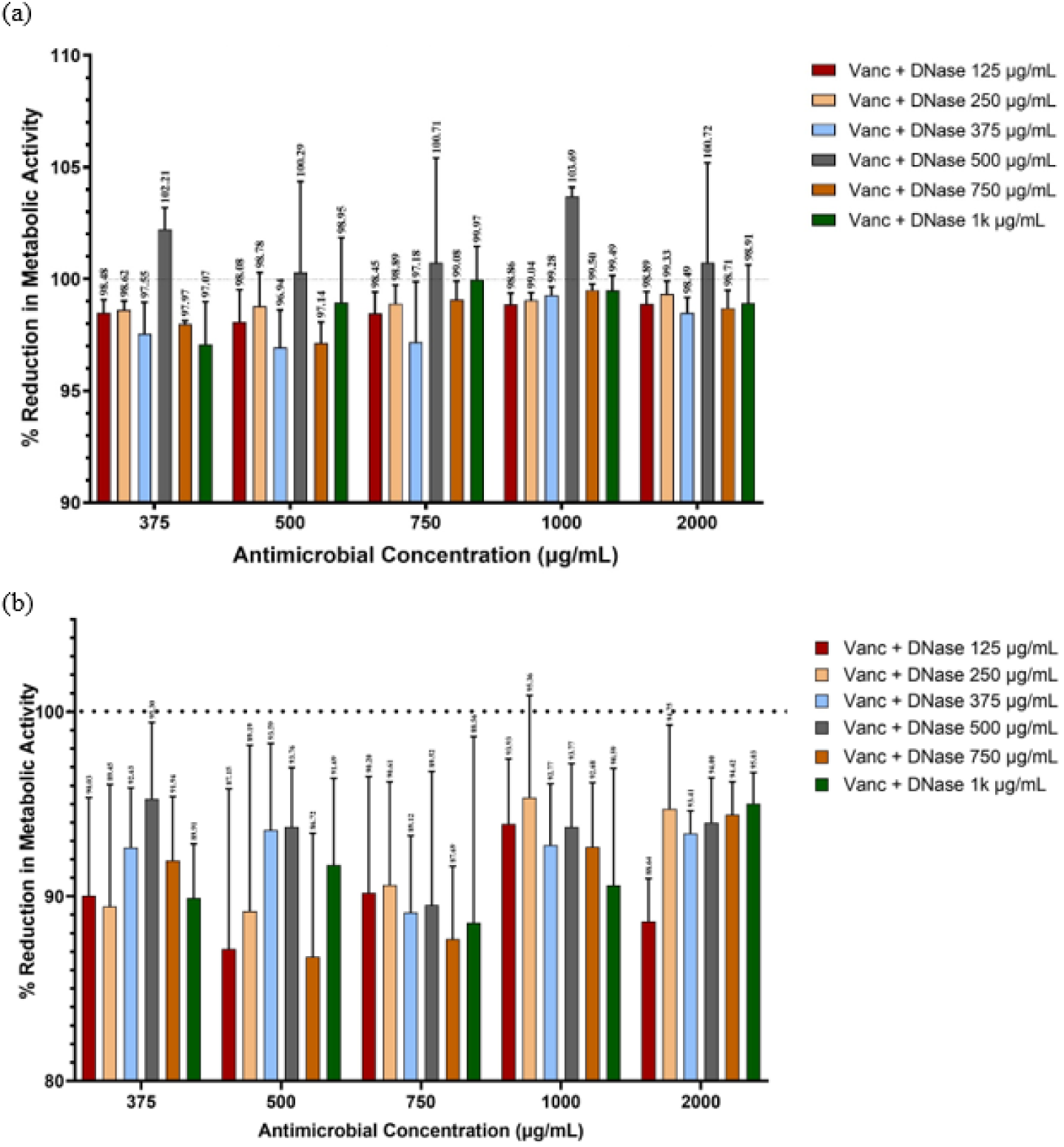
Effect of DNase I concentration and Vancomycin Combination on antimicrobial susceptibility of MRSA strains BAA-41 (a) and 33592 (b). Both strains of MRSA biofilm were grown for 72 hours in BHI broth supplemented with 1% glucose before antibiotic exposure for 24 hours. Antimicrobial susceptibility was determined based on reduction of metabolic activity determined with an XTT assay.

### 1.3.5 Determining the Presence of Persister Cells

Despite the promising results from the XTT assay, this marker tracks metabolic activity only. It is possible that the XTT assay will not account for the presence of low metabolically active persister cells. Also, the XTT assay in the biofilm model used here were normalized based on the untreated biofilm controls and in some samples, we observed more than a 100% reduction in metabolic activity. To address these potential weaknesses of the XTT screening assay, a subculture assay was developed that would allow the antibiotic treated biofilms to be subsequently grown for seven days in growth media without antibiotic challenge. The extended timeframe allowed for any remaining viable bacterial cells, persister or otherwise, to repopulate into the media and produce a turbid culture suggesting bacterial growth. The results from the experiment are presented in Table 2. It was observed that 1000µg/mL of vancomycin was generally insufficient to eradicate the MRSA strains despite the addition of DNase. Although little differences were seen in the XTT assay results performed under similar conditions (e.g. Figure 6(a)), the subculture assay for persister growth showed that higher vancomycin concentrations (2000 µg/mL) were crucial for consistently eradicating the biofilm.

**Table 2.**
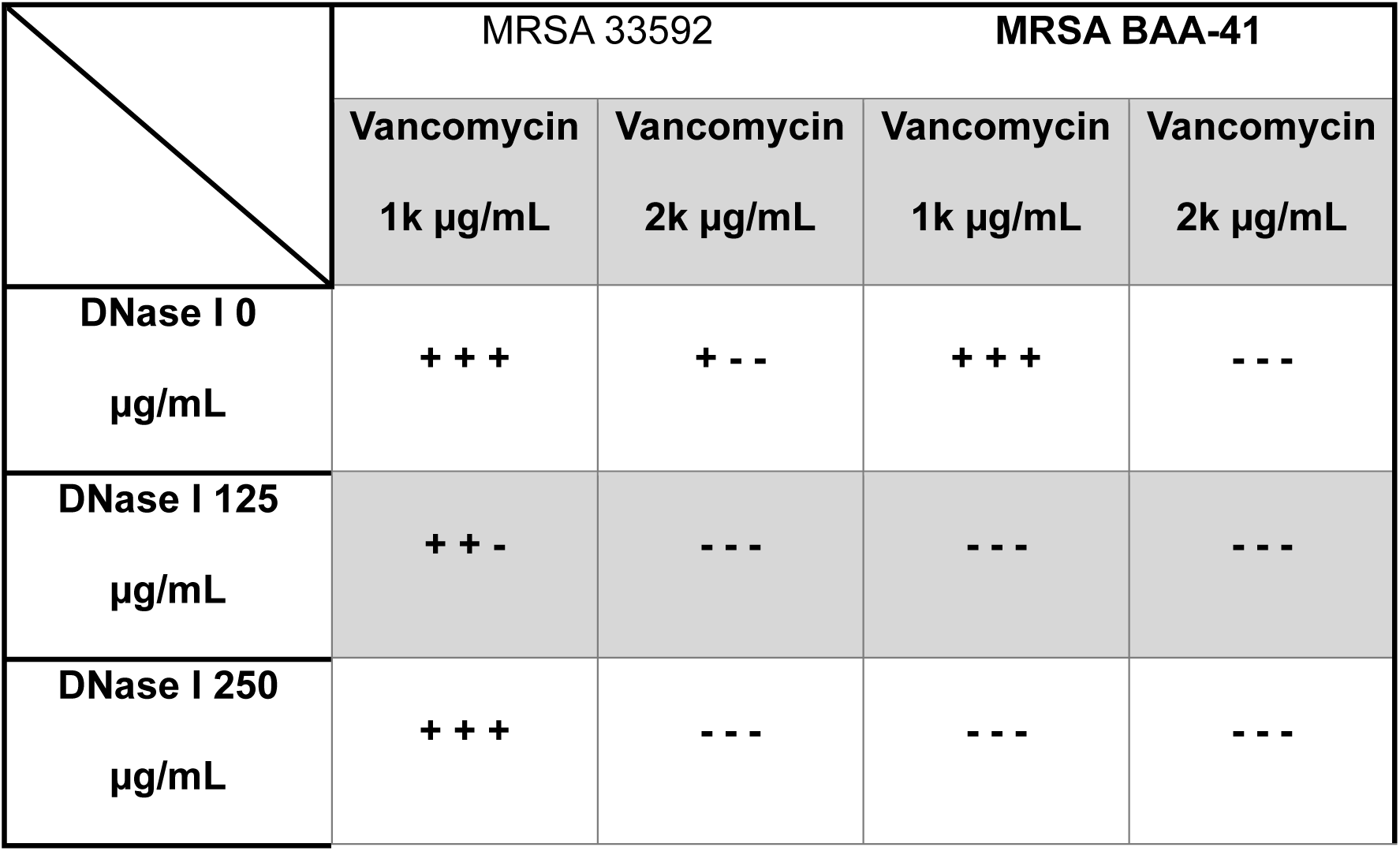
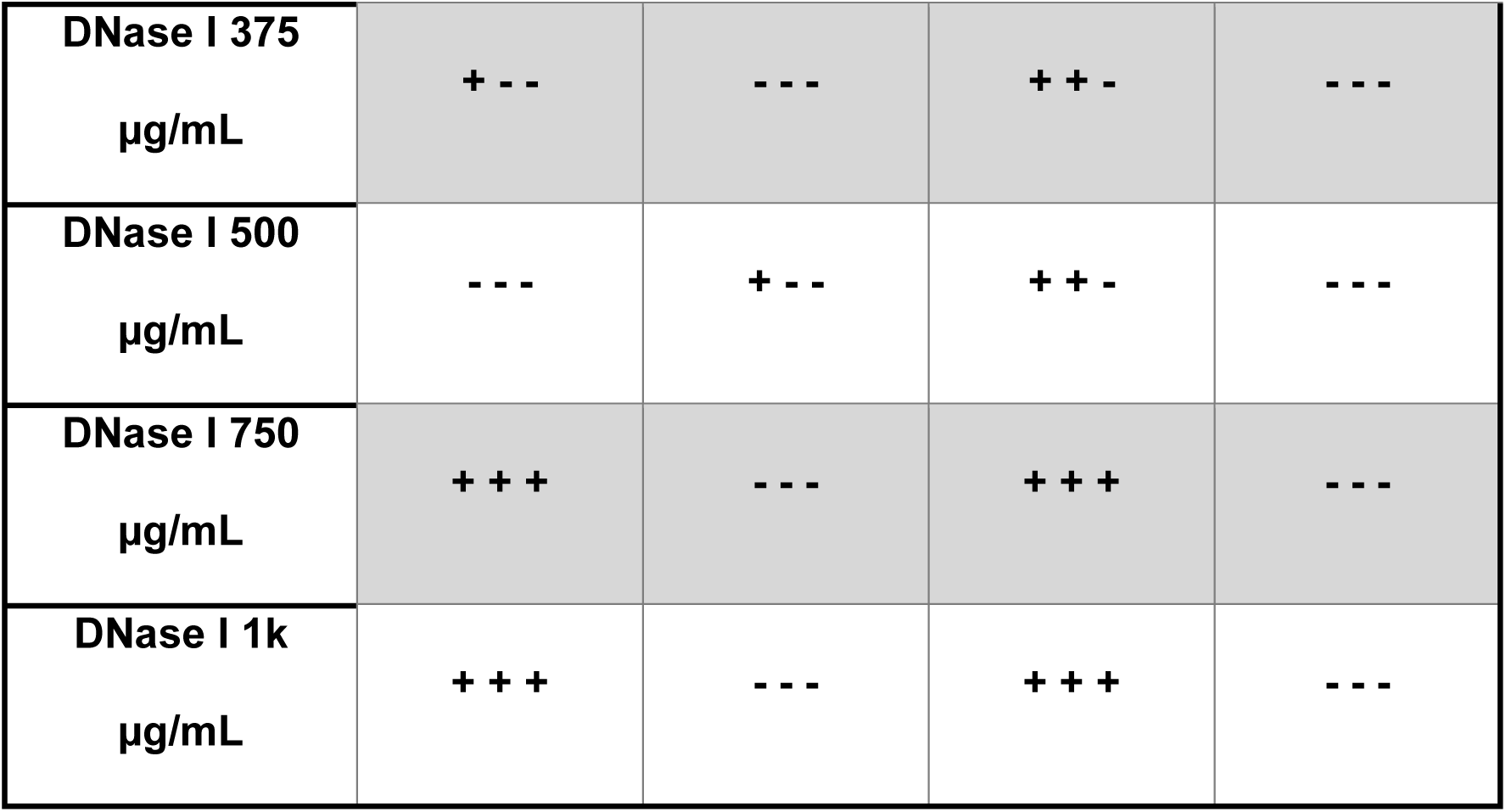
Culture results for MRSA strains 33592 and BAA-41. Samples that were positive for bacterial growth after 7 days are signified by a plus (+) sign, and samples negative for bacterial growth are signified by a negative (-) sign. For each condition, concentration of vancomycin and DNase, tested, three samples were analyzed.

### 1.3.6 Antimicrobial Diffusion in Biofilms

Since DNase by itself does not have bactericidal properties against the two strains of MRSA tested, the mechanisms of the improved antimicrobial performance of the DNase and vancomycin combination was further explored. One explanation could be that the inclusion of DNase improved the diffusion of vancomycin through the biofilm. To test this theory, a biofilm diffusion assay was utilized and the outcome for the experiment can be seen in Figure 7. The figure shows that the biofilms that were treated with both vancomycin and DNase saw significantly increased diffusion of vancomycin through the biofilm compared to the biofilms that were only treated with vancomycin. Statistical analysis showed the comparison between vancomycin and the co-delivered combination containing vancomycin and DNase was significantly different at all three timepoints for the BAA-41 strain with p-values of 0.029, 0.012, and 0.011 for 1, 2 and 3 hours respectively. The comparisons for the 33592 strain also resulted in significance with p-values of 0.048, 0.026, and 0.010 for 1, 2, and 3 hours respectively. Additionally, when comparing the apparent diffusivity coefficients as seen in Figure 8, both drug and enzyme combinations were statistically significant when compared to the antibiotic alone in the two MRSA strains tested (p-value < 0.0001).

**Figure 7.**
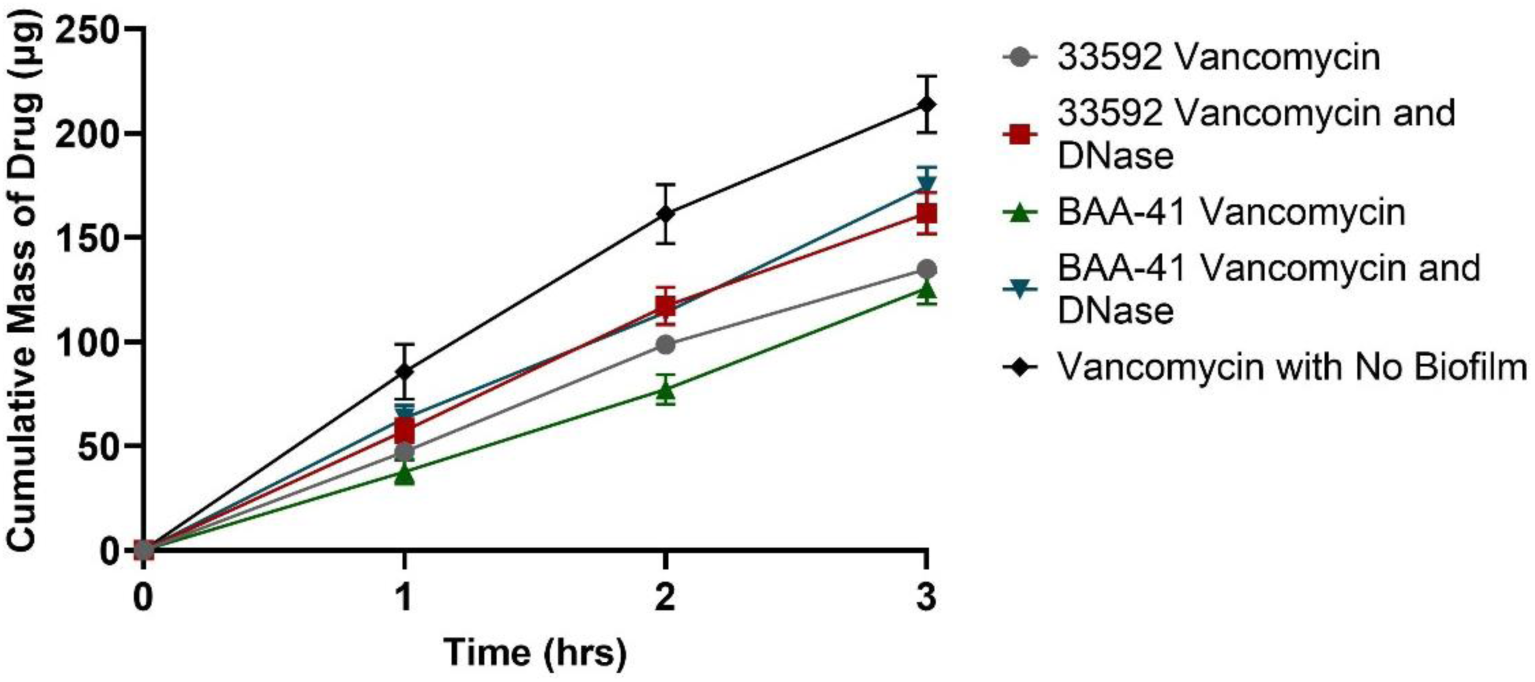
Effect of Vancomycin Diffusion for Vancomycin and DNase I Combination (a). Two different strains of biofilm, 33592 and BAA-41, were tested and compared to vancomycin without a biofilm as a control. 6.5mm Transwell® inserts with a 0.4 µm pore size were used to determine the diffusion of vancomycin through the biofilm. The inclusion of DNase resulted in improved diffusion of vancomycin but only the BAA-41 strain resulted in statistically significant improvement.

**Figure 8.**
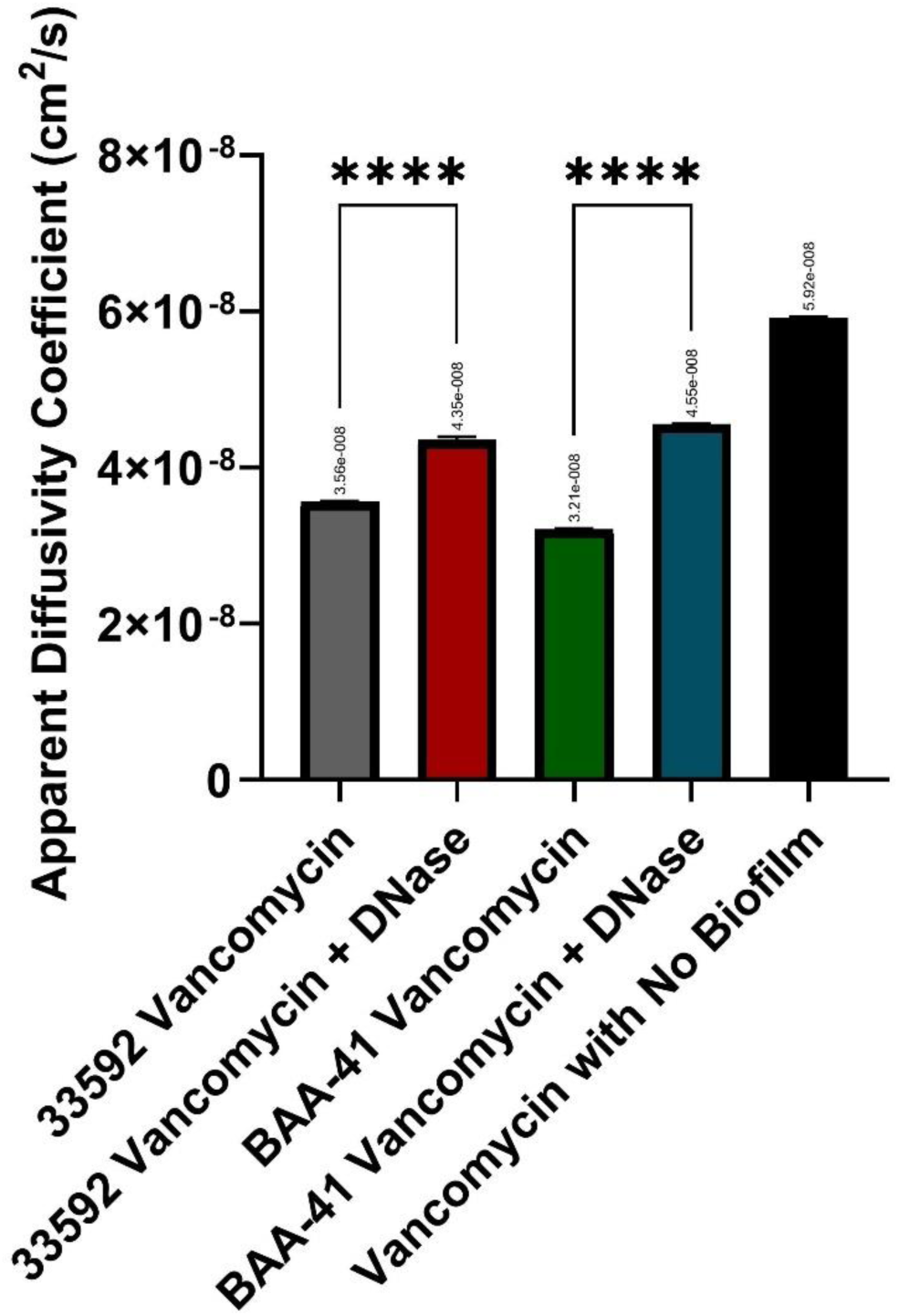
Apparent Diffusivity Coefficient. The diffusivity coefficient was calculated based on the data presented in Figure 7. The thickness, distance of diffusion, was estimated based on literature (42). The asterisks (****) denote statistical significance where in this case the p-value for the comparison between vancomycin and vancomycin + DNase combination was less than 0.0001 for both MRSA strains tested.

## 1.4 Discussion

### 1.4.1 Co-delivery of Antibiotics and Dispersal Enzymes Improves Antimicrobial Efficacy

The minimum biofilm eradication concentration (MBEC) was determined for the three antimicrobials identified. The antimicrobial performance against the biofilm was determined using an XTT assay which is a measurement of the metabolic activity of the bacterial cells within the biofilm. The results showed that over a 24-hour period, no single antibiotic was able to eradicate the biofilm, as determined by the reduction in metabolic activity. This is despite the concentrations tested (2000 µg/mL) being greater than levels achieved using common systemic routes of administration (2.4 – 9.6 µg/mL (17)). From Figure 3, it can be seen that MRSA strain 33592 was the more difficult strain to reduce metabolic activity. The exact mechanism for why strain 33592 has a lower antimicrobial susceptibility is not fully understood at this time. Figure 1 indicates that strain 33592 was a large biofilm biomass, but bulk diffusion, as shown in Figure 7 in the case of vancomycin demonstrates that strain 33592 does not provide a greater barrier to the permeation of the antimicrobial. Diffusion on the microscale may be different to that on the bulk scale (31, 43, 44). Another possible explanation is the differences in biofilm composition or regulatory and defense mechanisms which would allow for strain 33592 to be more robust to antimicrobial challenge. Another interesting observation is that clinical literature suggests that linezolid is superior to vancomycin when used to treat MRSA pneumonia infections (8). Since only two strains of MRSA were tested here it is not possible to extrapolate the results to all strains, but statistically there is not a difference between linezolid and vancomycin when equal concentrations are exposed to the biofilms under similar *in vitro* conditions. Clinical studies such as the Zephyr study (8) provide valuable information, but systemically administered antibiotics are limited by the pharmacokinetic and distribution properties of the drug. The clinical performance of a systemically delivered antimicrobial may not directly relate to its success, or lack thereof, of the same locally delivered antimicrobial. An interesting observation in regard to the performance of tigecycline can be seen in Figure 3 where the increasing concentrations of the antimicrobial see a decreasing antimicrobial susceptibility of the biofilm. Literature has shown increasing concentrations of tigecycline are more susceptible to accelerated degradation resulting in a lower amount of viable antimicrobial exposed to the biofilm (45). This conclusion was supported by the observation that the higher concentrations of tigecycline resulted in a darker, almost black color, of the samples at the end of the timepoints tested which is an indicator of tigecycline degradation (46, 47).

Following the inability for any of the three antibiotics to eradicate the biofilm after 24 hours, the two best performing antimicrobials, vancomycin and tigecycline, were supplemented with either DNase or proteinase K. Our working hypothesis was that the inclusion of DNase or proteinase K would disrupt the biofilm by reducing the amount of extracellular DNA and/or proteins present in the biofilm structure allowing improved antibacterial action. The results shown in Figures 4 and 5 support the idea that the inclusion of DNase or proteinase K weakens the biofilm resulting in the improved performance of the antimicrobials tested. It is important to note that the inclusion of DNase or proteinase K resulted in statistically significant improvements in antimicrobial performance, but the remaining biofilm was still viable following culturing suggesting that statistical significance alone is not sufficient in determining treatment success when considering *in vitro* biofilm models. Even with the improved performance with the inclusion of the dispersal agents, the biofilm needed to be exposed to high concentrations of antimicrobial (2000 µg/mL) and dispersal agent combination for 24 hours for strain BAA-41 and 72 hours for strain 33592 to eradicate the biofilm with a 100% reduction in metabolic activity. The high concentration of antimicrobial greatly exceeds what is typically seen with systemically delivered formulations (17), but other groups have found that inhaled therapies are able to achieve the high required doses in the lung while maintaining safe systemic levels (48). To mimic dosing frequency and limit concerns related to drug degradation, the drug combination was replaced every 12 hours. Despite DNase and proteinase K having different targets within the biofilm, both greatly improved the performance of antimicrobials especially in the case of vancomycin suggesting that both proteins and extracellular DNA are susceptible targets for therapeutic agents. Neither DNase nor proteinase K were able to have a pronounced effect on the biofilm without being paired with an antimicrobial which is supported by others (29, 49–52). Following performance of the antimicrobial and dispersal agent combinations, it was determined that the combination of vancomycin and DNase offered the best path to successfully eradicating biofilms. Across the two strains tested, the vancomycin and DNase combination demonstrated excellent ability for eradicating the biofilm determined by the XTT assay. Since DNase is already in an inhaled approved FDA product, Pulmozyme, it possesses an established safety profile in the route of administration that is being pursued. Also, the co-delivery of the of vancomycin and DNase resulted in an improved, albeit non-signifincant, killing of the biofilm compared to a staggered delivery of DNase followed by vancomycin, as one may seen in the clinically setting. The staggered delivery did offer significant improvement over vancomycin alone when exposed to the 33592 biofilm for 12 hours when tested. It is the opinion of the authors that this difference between the staggered and co-delivery of vancomycin and DNase would result in differences in clinical treatment outcomes. Additional concentrations of DNase in combination with vancomycin were tested to determine the potential formulation space that is available. The results in Figure 6 show that concentrations ranging from 125-1000 µg/mL had no significant effect on the performance the antimicrobial and dispersal agent combination. The flexibility in DNase concentration is important when it comes to formulating an inhaled drug product since the overall powder load should be limited as much as possible (53).

### 1.4.2 Codelivery leads to eradication of persister cells

Although XTT assays are widely used to determine the antimicrobial susceptibility of *in vitro* biofilms, one severe limitation they possess is that they rely on the metabolic activity of the bacterial cells. A measurement of metabolic activity is not a good assay for the presence of persister cells due to their low or absent metabolic activity. A broth subculture assay was performed after antimicrobial exposure to determine the presence of persister cells. The subculture samples were monitored over a 7-day period to check for bacterial growth determined by the turbidity of the culture. Both positive and negative controls of untreated biofilm and sterile media were used to ensure the accuracy of the assay. Despite reaching the 100% reduction in metabolic activity goal, an overwhelming majority of the samples tested that contained 1000 µg/mL of vancomycin, as seen in TABLE 2, were positive for bacterial growth in both MRSA strains tested (i.e. persisters were present). This table shows that the samples containing 2000 µg/mL were sufficient to eradicate the biofilm under the tested conditions (two samples were believed to be false positives due to contamination during testing). One concern with the use of DNase as the dispersal agent, was that varying strains of MRSA will have different concentrations of extracellular DNA within the biofilm. Although the composition of the two MRSA strains tested was not analyzed, this scenario was tested by using only vancomycin to determine if the antimicrobial alone was sufficient to eradicate the biofilm. As seen in TABLE 2, it was determined that vancomycin alone was sufficient to eradicate the biofilm if given an appropriate exposure time. In the absence of an immune response, the results show that vancomycin with and without the addition of DNase was sufficient to eradicate the robust mature biofilms that were formed.

### 1.4.3 Codelivery results in improved antimicrobial diffusion

Following the positive results of the vancomycin and DNase drug combination, we wanted to determine the underlying mechanism that resulted in the improved performance of the combination. Initially, the possibility that vancomycin was seeing reduced diffusion through the biofilm was investigated (27, 30, 54). To test this theory, a diffusion assay was utilized which showed that the inclusion of DNase significantly improved the diffusion of vancomycin through the two MRSA strains studied. A possible explanation for the improved diffusion could be that the structure of the biofilm was compromised as extracellular DNA was degraded by DNase leading to improved diffusion of vancomycin. Another possible explanation is that extracellular DNA has been shown to chelate certain antibiotics such as vancomycin (42, 55). The combination of DNase and vancomycin may have allowed for more vancomycin to be available to treat the biofilm by reducing the amount of antimicrobial chelated by the extracellular DNA present in the biofilm structure. An additional benefit of including DNase in the drug combination is that extracellular DNA has been proven to play a critical role in horizontal gene transfer which is one of the critical components of resistance development within biofilms (29, 42, 55). The impact of antimicrobial resistance can be mitigated by the eradication of all the cells within biofilms. The inclusion of DNase has the potential to prevent horizontal gene transfer reducing the chance of resistance development. The theory of preventing resistance development was not studied as part of this work so further analysis needs to be undertaken to confirm the validity of this possible approach.

### 1.4.4 Rigorous *in vitro* biofilm model development

A rigorous *in vitro* biofilm model was developed by first analyzing the potential impact that growth conditions can have on a biofilm. First, the impact of growth conditions on biofilm biomass was determined using a crystal violet assay. In total, six different growth medias were tested over three different time periods for two different MRSA strains. Although MRSA strain 33592 produced a larger biofilm compared to BAA-41, a 3-day growth with BHI supplemented with 1% glucose resulted in the largest biofilm without sacrificing reproducibility. Identifying a robust biofilm was important because under these *in vitro* conditions the immune response was not considered; so, it was important to determine whether a biofilm could successfully be treated with antibiotics in what essentially amounts to a worst-case treatment scenario. One interesting thing to note is that despite the lack of statistical significance of biofilm biomass between the two MRSA strains tested using a 3-day growth period in BHI + 1% glucose, the antimicrobial performance under the same growth conditions was considerably different which suggests that statistical significance, or lack thereof, may not always translate to potential clinical significance.

In addition to the biofilm biomass, another important metric that was determined was the impact of the growth conditions on the number of colony forming units within the biofilms tested. Since crystal violet assays do not account for the viability of the cells within the biofilms tested, a drop-plate method was used on biofilms grown under similar conditions to the biomass testing. Although biomass testing showed discernable graphical differences between the two strains of MRSA tested at various growth conditions, the number of colony forming units within the biofilms were more similar. However, upon statistical analysis using Tukey’s multiple comparison, there was more difference between the samples tested than what was seen during biomass analysis. It is interesting to note that although BAA-41 grown for 3-days in BHI + 1% glucose resulted in a biofilm that was about 40% smaller than 33592 under the same conditions, the number of colony forming units for each strain was almost identical, with no statistical difference, with a log CFU/mL of 5.999 for BAA-41 and 6.026 for 33592. One of the major concerns with an *in vitro* test is its applicability to *in vivo* conditions especially in the case of antimicrobial susceptibility testing where the CFU count can be shown to have a large impact. The log of CFU/mL in a clinical setting can vary widely, but typically the median falls between 5.8 and 6.6 (56–58). The biofilm growth with BHI + 1% glucose at 3-days, identified as the best performer in terms of biomass, fell within that expected clinical range. The combination of the robust biofilm formed populated with a clinically expected population of bacteria suggests that biofilms grown for three days in BHI + 1% glucose were a suitable model for testing antimicrobial susceptibility. Another observation made was that the variability, as seen by the error bars in Figure 2, within sample groups was much smaller for the number of CFUs compared to biomass suggesting that biomass may be more sensitive to stimuli than the number of CFUs within the biofilm. The consistency, as a factor of relative standard deviation, in the number of CFUs is not incredibly surprising due to the high amount of regulation within a biofilm, but it does raise questions about why the less consistent growth of the biomass had apparently little effect on the number of CFUs under similar circumstances. This data would suggest that the number of CFUs is more heavily regulated and robust to stimuli than the biomass of the biofilms themselves; however, further analysis would be needed to confirm this assumption. Understanding the intricacies of biofilm growth is important for the development and improvement of therapeutic treatments.

Following the determination of the best growth conditions for the proposed set of experiments, MIC testing was undertaken to act as a screening tool for potential antibiotics. Six antimicrobials were chosen with three (vancomycin, linezolid, and clindamycin) recommended in clinical guidelines and an additional three (tigecycline, tobramycin, and niclosamide) that have had varying levels of success in literature. The results showed that vancomycin, tigecycline, and linezolid performed the best with the remaining three antimicrobials having less than optimal impacts. Due to these results, it was determined that the three successful antimicrobials warranted further study in a mature biofilm model. It is important to note that the MIC values found for vancomycin were higher than what is typically reported which could be a factor of the MRSA strains tested. One interesting observation was that tigecycline vastly outperformed either vancomycin and linezolid in terms of MIC testing, but the same outcome was not seen in the MBEC testing suggesting that MIC analysis may not be a beneficial test when a biofilm infection is present despite the widespread use of MIC testing in the development of and use of antimicrobials. In the two strains of MRSA tested, tigecycline outperformed the other two antimicrobials tested in the 33592-strain shown in Figure 3(b), but in the BAA-41 strain tigecycline was outperformed despite vancomycin having an MIC that was 8-fold higher than tigecycline. It would be expected that the antimicrobial with the lower MIC would result in a better performance against an *in vitro* biofilm model, but that outcome did not come to fruition which is further supported by the results showing the inclusion of the dispersal agents in the form of DNase and proteinase K in Figures 4 and 5.

### 1.4.5 Limitations of the study

This study uses an *in vitro* model that does not fully account for the complexities of a clinical biofilm infection. Although the *in vitro* model offers a simplified approach since it does not account for the immune response among numerous other variables, a robust biofilm was still able to be grown that proved difficult to treat. The biofilms were grown under essentially ideal conditions since there was plenty of nutrients present as well as no challenge except for the antimicrobials used. It is likely possible that in a clinical setting the concentration of antimicrobial would be less than what was presented here due to changes in available nutrients or challenges made by the immune system. Another limitation is that the antibiotic concentrations that were tested are not feasible through typical systemic routes of administration that are used clinically. However, other routes of administration are available such as pulmonary delivery, which has been proven to deliver high doses in the Tobi® Podhaler® and AeroVanc™ previously referenced.

## 1.5 Materials and Methods

### 1.5.1 Materials and Bacterial Strains

Two methicillin resistant *Staphylococcus aureus* strains (ATCC #: BAA-41 and 33592) were obtained from ATCC (Manassas, VA). Brain heart infusion broth (BHI) from VWR (Radnor, PA) and tryptic soy broth (TSB) from Becton Dickinson (Franklin Lakes, NJ). The antibiotics linezolid, clindamycin, and tigecycline were purchased from Ambeed (Arlington Heights, IL). Vancomycin was purchased from VWR. Niclosamide was purchased from Shenzhen Neconn Pharmatechs Ltd (Shenzhen, China); and tobramycin was purchased from Letco Medical (Decatur, AL). Proteinase K and DNase I were acquired from MP Biomedicals (Santa Ana, CA) and Roche Diagnostics (Mannheim Germany), respectively. XTT Cell Viability Assay kit was purchased from Biotium (Freemont, CA).

### 1.5.2 Preparation of Inoculum

A single colony of each MRSA (ATCC #: BAA-41 and 33952) strain was removed from tryptic soy agar plates and added to culture tubes containing 3 mL of either TSB or BHI media. The culture tubes were sonicated in a water bath for 1 minute before being placed in a 37°C incubator overnight (20-24hrs).

### 1.5.3 Biofilm Biomass by Crystal Violet Assay

Biofilm biomass was determined using a crystal violet assay similar to a protocol developed by Sugimoto et al (29). To briefly summarize the assay, following the preparation of the overnight inoculum, two 3 mL overnight inoculum cultures (6 mL total) were diluted with 100 mL of BHI or TSB media supplemented with either 1% (w/v) glucose or 2 or 3% (w/v) sodium chloride. The growth media compositions tested were BHI, BHI + 1% glucose, BHI + 3% sodium chloride, TSB, TSB + 1% glucose, and TSB + 2% sodium chloride). 200 µL of the suspension was added to 96-well flat bottom polystyrene tissue culture plates (Corning, NY) and incubated at 37°C for 1, 3, or 5 days. After the designated growth period, the wells were washed twice with 200 µL of sterile phosphate buffered saline (PBS) before 200 µL of 0.05% Crystal Violet (CV) was added to the wells. The crystal violet stain was applied for 5 minutes before the biofilms were washed twice with PBS before biofilm biomass was quantified by measuring the absorbance at 595 nm with a Tecan infinite M200 (Mannedorf, Switzerland) microplate reader.

### 1.5.4 Counting of Colony Forming Units

The number of colony forming units (CFUs) was determined using a drop plate protocol modified from an article published by Miles, Misra and Irwin (59). To summarize, following the same growth conditions outlined in the previous biofilm biomass section, the wells containing the biofilms were washed with sterile BHI or TSB media. Afterwards, the biofilm was removed from the well by using a sterile pipette tip to scrap the bottom and walls of the well to remove any bacteria. The bacteria on the pipette were transferred to culture tubes containing 3 mL of either BHI or TSB media before being placed in a water bath sonicator for 1 min. The culture tubes were vortexed on a medium setting for 30 seconds to 1 minute before 10x serial dilutions were performed. 20 µL of each dilution was added to tryptic soy agar plates which were incubated at 37°C for 24 hrs before the number of colonies was counted.

### 1.5.5 Minimum Inhibitory Concentration (MIC)

Minimum inhibitory concentration (MIC) was determined based on antimicrobial susceptibility testing standards published by the Clinical and Laboratory Standards Institute (CLSI) (40). To briefly describe the method, six antimicrobials (linezolid, clindamycin, vancomycin, niclosamide, tobramycin, and tigecycline) were tested at concentrations ranging from 0.5-64 µg/mL in 2x serial dilutions. The bacterial inoculum (either BAA-41 or 33592) used was prepared as shown earlier but was diluted to 0.5 McFarland standard before 10 µL of inoculum was added to a flat-bottom 96-well plate. 100 µL of antibiotic was added to each well of the 96-well plate with the bacterial inoculum before 24-hrs of incubation at 37°C. The MIC was determined for the lowest concentration of antimicrobial which did not produce a turbid sample.

### 1.5.6 Minimum Biofilm Eradication Concentration (MBEC)

The minimum biofilm eradication concentration (MBEC) was determined using MRSA biofilms (either BAA-41 or 33592 strains) that were prepared similarly as shown in previous sections. To summarize, the overnight inoculum was added to 100 mL of BHI supplemented with 1% (w/v) glucose before 200 µL of suspension were added to flat-bottom 96-well plates. The biofilms were incubated for three days at 37°C. Following incubation, the biofilms were washed once with sterile BHI growth media before 200 µL of one of three antimicrobials (vancomycin, linezolid, or tigecycline) were added at concentrations ranging from 32-2000 µg/mL in 2x serial dilutions for antibiotic exposure times ranging from 12 to 72 hrs. The antibiotic was applied in two manners, either a one-time initial dose or replacing the antibiotic every 12 hrs. Following the appropriate antibiotic exposure time, the remaining antibiotic media was removed from the well and the biofilm was washed with 200 µL of sterile media. The preparation of the XTT assay followed the manufactures guidelines where 25 µL of the activating agent was mixed with 5 mL of XTT solution. 150 µL of sterile media was added to each sample as well as 75 µL of the XTT and activating agent solution. Following a three-hour incubation at 37°C, the absorbance was determined at 490nm using a Tecan infinite M200 microplate reader. The impact of passive dispersal agents, in the form of DNase I (at concentrations ranging from 125-1000 µg/mL) and Proteinase K at 100 µg/mL was determined when the passive dispersal agent was used in combination with either vancomycin or tigecycline with antibiotics exposure times ranging from 12 to 72 hrs. The MBEC was determined for samples that resulted in a 100% reduction in metabolic activity.

### 1.5.7 Subculture to detect persister cells and slow growing bacterial cells

Since XTT assays typically do not account for persister cells that inhabit biofilms, a method modified from Badha et al. was used (41). To briefly summarize the method, the biofilm was grown and treated with antibiotics, either solo or in combination with a passive dispersal agent, as already mentioned. Vancomycin (either solo or in combination with DNase I) was tested with vancomycin concentrations of 1000 and 2000 µg/mL and DNase concentrations ranging from 125 to 1000 µg/mL. BAA-41 biofilm samples were treated with antibiotics for 24-hrs replacing the antibiotic after 12-hrs while 33592 was exposed to the antibiotic for 72-hrs again replacing the antibiotic every 12-hrs. Following the treatment of the antibiotic, the wells containing the biofilm samples were washed once with 200 µL of sterile media. Following the removal of the wash media, 200 µL of media was added to the well, and a pipette tip was used to scrape the bottom and walls of the well to suspend any remaining bacterial cells in the media. The suspension was then transferred to culture tubes containing 2.8 mL of media which was sonicated in a water bath for 1 minute. The sonicated culture tubes were then incubated for 7 days. Following a week of observation, the presence of bacterial growth was determined by the turbidity of the sample, or lack thereof for negative samples.,. The MBEC for all cell populations within the biofilm was determined for the samples that resulted in negative subcultures with clear breakpoints for the various concentrations tested.

### 1.5.8 Antibiotic Diffusion through the biofilm

To determine the impact of DNase on the diffusion of vancomycin through the biofilm, a diffusion assay was modified from Gonzales et al. (60). To briefly summarize the method, 24-well Transwell® plates were used with a 0.4µm pore size membrane. The biofilm of the two MRSA strains was grown as previously mentioned over a 72-hour period. Following sufficient biofilm growth, 450 μL of deionized (DI) water were added to the basolateral side of the membrane and 400 μL of DNase and vancomycin were carefully added on top of the biofilm, apical side, without disturbing the biofilm. The biofilm was then incubated before samples were collected every hour over a three-hour period. The collected samples were analyzed using a high-performance liquid chromatography (HPLC) method developed by Wicha and Kloft (56). An Accucore® C-18 HPLC column (2.6μm, 100 x 2.1 mm, Thermo Fisher, Dreieich, Germany) was attached to a Dionex 3000 HPLC (Thermo Fisher) with UV detection at 240 nm controlled using the Chromeleon® software. Mobile phase A consisted of DI water with 0.1% v/v trifluoroacetic acid (TFA), and mobile phase B consisted of acetonitrile with 0.2% TFA. A gradient profile was used consisting of 15% mobile phase B for 2 minutes followed by 75% mobile phase B for 8 minutes, and finally 15% of mobile phase B for the final 2 minutes of the 14-minute run.

### 1.5.9 Statistical analysis

Statistical analysis was performed using GraphPad Prism (Version 10) statistical software to analyze the data using two-way analysis of variance (ANOVA) with the addition of a Tukey test for multiple comparisons unless otherwise noted. Data shown is presented as mean and standard deviation unless stated differently.

## 1.6 Conclusion

The eradication of mature MRSA biofilms requires high concentrations of antimicrobials that greatly exceed the MIC by multiple orders of magnitude. Additionally, a single antibiotic is unlikely to lead to the eradication of an *in vitro* biofilm. The incorporation of biofilm disruptors such as DNase I or Proteinase K greatly improves the improves the antimicrobial performance of vancomycin or linezolid. However, the safety profile for Proteinase K *in vivo* has not been well established. Alternatively, DNase I has a well-established safety profile due to the aerosolized Pulmozyme formulation for cystic fibrosis patients, which reduces the viscosity of mucus in the lung. The combination of DNase I and vancomycin was able to eradicate mature MRSA biofilms, including persister cells, within 72 hours of antimicrobial exposure. DNase I was able to improve the diffusion of vancomycin into the biofilm likely accounting for the improved performance. The combination of DNase I and vancomycin provides a potential solution for the treatment of MRSA biofilm lung infections.

## 1.7 Acknowledgments

This research received no specific grant from any funding agency in the public, commercial, or not-for-profit sectors.

HDCS, an author of this paper, consults for and has equity ownership in Via Therapeutics, Nob Hill Therapeutics and Cloxero Therapeutics on inhaled product development. The terms of this arrangement have been reviewed and approved by the University of Texas at Austin in accordance with its policy on objectivity in research.

RCM, an author of this paper, has equity ownership in Cloxero Therapeutics, a company focused on the delivery of antimicrobial agents locally to the lung for the treatment of bacterial lung infections. The terms of this arrangement have been reviewed and approved by the University of Texas at Austin in accordance with its policy on objectivity in research.

